# Interpretable Convolution Methods for Learning Genomic Sequence Motifs

**DOI:** 10.1101/411934

**Authors:** MS Ploenzke, RA Irizarry

## Abstract

The first-layer filters employed in convolutional neural networks tend to learn, or extract, spatial features from the data. Within their application to genomic sequence data, these learned features are often visualized and interpreted by converting them to sequence logos; an information-based representation of the consensus nucleotide motif. The process to obtain such motifs, however, is done through post-training procedures which often discard the filter weights themselves and instead rely upon finding those sequences maximally correlated with the given filter. Moreover, the filters collectively learn motifs with high redundancy, often simply shifted representations of the same sequence. We propose a schema to learn sequence motifs directly through weight constraints and transformations such that the individual weights comprising the filter are directly interpretable as either position weight matrices (PWMs) or information gain matrices (IGMs). We additionally leverage regularization to encourage learning highly-representative motifs with low inter-filter redundancy. Through learning PWMs and IGMs directly we present preliminary results showcasing how our method is capable of incorporating previously-annotated database motifs along with learning motifs *de novo* and then outline a pipeline for how these tools may be used jointly in a data application.

## Introduction

Applications of deep learning methods have become ubiquitous over recent years due primarily to excellent predictive accuracy and user-friendly implementations. One such application has been to nucleotide sequence data, namely data arising in the field of genomics, in which the convolutional neural network (CNN) has enjoyed particular success. The convolutional layers composing a CNN work by extracting and scoring local patches of the input data by computing the cross-correlation between all nucleotide subsequences in the observation and each filter. These feature scores are then passed through any number of subsequent weightings (so-called dense or fully-connected layers) and used to output a final predictive value or values, as in the case of a multi-dimensional output. For example, one of the earliest CNNs trained on genomic data, DeepSea, predicted with high accuracy a 919-dimensional output array with each entry representing the presence/absence of a specific chromatin feature [29]. DeepBind, developed near the same time as DeepSea, further demonstrated the utility of training CNNs on genomic data by showcasing how the first-layer convolutional filters tend to learn relevant sequence *motifs* [2]. This latter finding highlighted, within the application to genomic data, the potential for illuminating the black box that deep models are typically considered; namely it sparked interest in developing computational methods for both incorporating known biological structure into the models [1, 7, 16, 23] as well as interpreting the learned model knowledge [12,18,22].

Much progress has been made to improve predictive accuracy since these pioneering manuscripts however the process proposed by [2] to infer sequence motifs from convolutional filters remains largely unchanged. Specifically, each trained filter is convolved over input test set observations to produce a vector of activation values per filter per observation. High scoring activation values above some threshold are identified and the subsequence giving rise to each value is extracted. All extracted subsequences are stacked together per filter and used to compute a position frequency matrix (PFM). The PFM for filter *j* is a 4 × *L*_*j*_ matrix in which the rows represent nucleotide (A, C, G, T) and columns represent position. *L*_*j*_ is generally on the order of 8-18bp. The columns may be subsequently normalized by their sum to yield a *position probability matrix* (PPM), and then converted into a *position weight matrix* (PWM) by computing, for each element *ω*_*n*_ in the PPM, *log*_2_(*ω*_*n*_) – *log*_2_(*b*_*n*_) where *b*_*n*_ represents the background probability for the nucleotide *n*. The PWM is often visualized as the so-called sequence logo [20], which is computed by multiplying each entry in the PPM by the column-wise sum of the expected self-information gain (i.e. the Hadamard product of the PPM and PWM). Sequence-logo motifs constructed and visualized in this manner are shown in the bottom rows of Fig. 1A-D. We refer to the matrix of values denoting the heights of the nucleotides as the *information gain* m*atrix* (IGM) of the PWM. A worked example converting a PFM to an IGM is provided in the supplementary materials.

**Figure 1.**
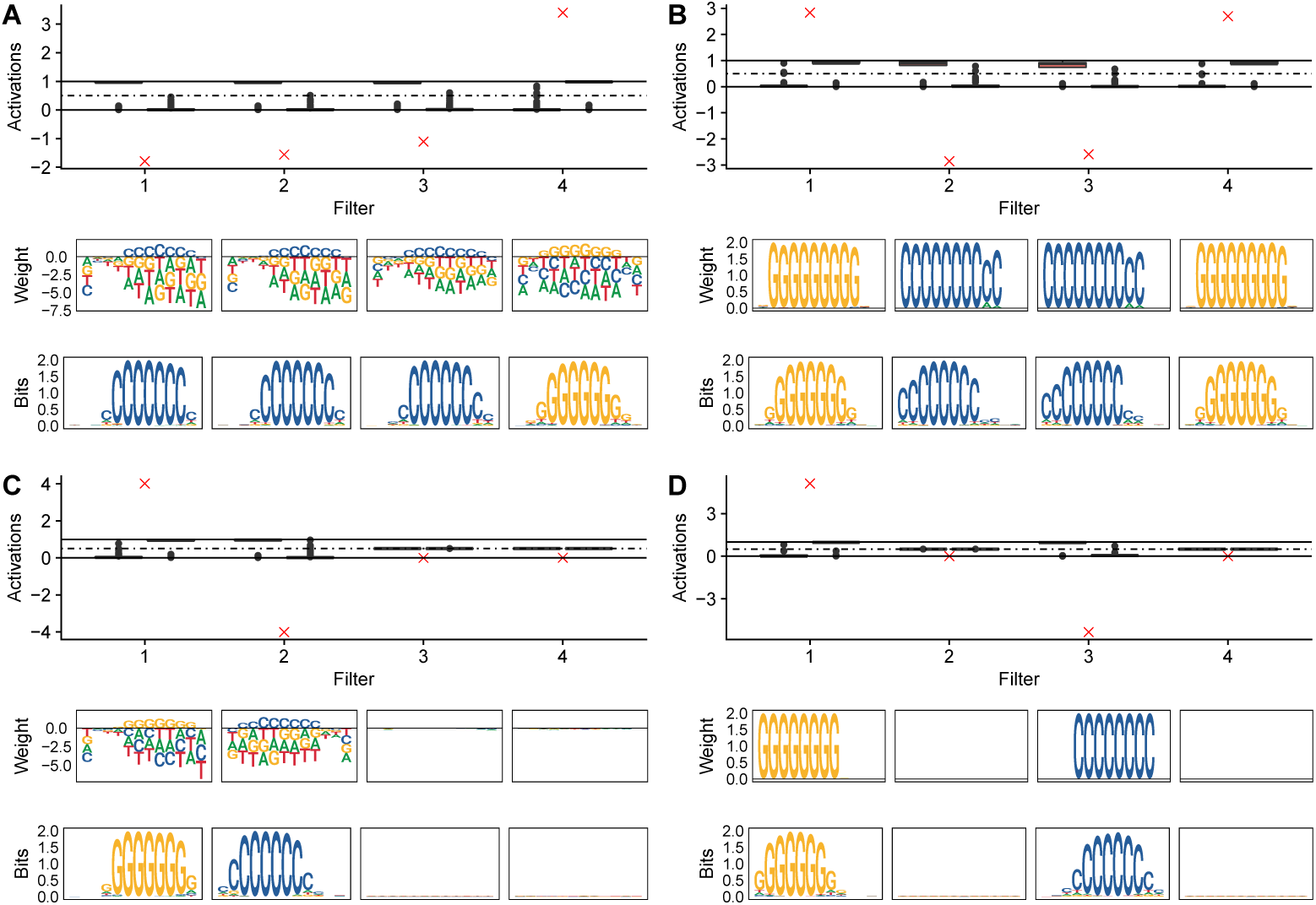
Effect of weight constraints and regularization on learning motifs. Box plots show first-layer convolutional filter activations post sigmoid transformation by class label. *Y* = 0 sequences containing the *CCCCCCCC* motif achieve large activations with the *C*-motif filters (filters 1, 2, 3), *Y* = 1 sequences containing the *GGGGGGGG* motif achieve large activations with the *G*-motif filter (filter 4). Red ×‘s indicate the associated 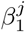 coefficients (effect size estimates). **A.** Unconstrained filters (middle row) within an unregularized model (top row) learn redundant sequence motifs and require test set observations for motif interpretation (bottom row). **B.** Unregularized filters constrained to represent valid IGMs do not require test set observations and are directly interpretable as sequence-logo motifs. Filter redundancy remains. **C.** Filter regularization discourages learning redundant features. **D.** Constrained filters within a regularized model learn distinct sequence motifs directly with no need for post-hoc interpretation procedures.

Under the standard CNN framework there are no restrictions on the values of the weights comprising each convolutional filter to any range. In addition, the visualization procedure requires the analyst to select the threshold for extracting high activation values and is dependent upon the input observations themselves. As illustrated in panels A and B of Fig. 1 however, filters tend to learn redundant and highly-correlated features. Given the non-identifiability of the many weights in a typical CNN it is no wonder why the learned features are so correlated. One recently-proposed technique specifically developed to remedy the issue of correlated filters is to include all possible circular shifts of the filter when convolving it with the sequence [3]. Such a procedure increases the computation time and memory footprint as every convolutional operation now requires all possible spins of the filter and also requires input observations to interpret what has been learned. An alternative approach somewhat abandons the notion of filter interpretability as a sequence motif and instead takes a reverse approach via back-propagating the activation values, in effect addressing which sequences are most important for model classification for a given observation relative to some reference observation [22]. A third approach involves solving a reformulated optimization problem which seeks to find the single consensus sequence maximally activating the model [12,18]. None of these techniques simultaneously address the issues of redundancy and interpretability and, moreover, they require input observations or a post-hoc optimization procedure to infer the learned motif.

We propose to directly learn the sequence motifs such that interpretation of the convolutional filters is not reliant upon test set observations and the weights comprising the filter are directly interpretable as information gain or position weights, both of which may be easily visualized as sequence logos. We simultaneously address the issue of filter redundancy along with interpretability by incorporating weight constraints and regularization techniques. The weight constraints limit the range of the individual filter weights to restrict their values as to be directly interpretable as IGMs or PWMs while the regularization scheme encourages learning non-redundant motifs. Under such a framework previously-annotated database motifs either in the form of PWMs or IGMs, such as those available from JASPAR [10], may be used to initialize the convolutional filters in the model and subsequently held constant or updated during training. In section 2 we provide a brief introduction to the notation that will be used before detailing the method through a toy simulation. Section 3 showcases results for a more realistic simulation study as well as a data example using ChIP-seq peaks from the ENCODE Consortium [6]. Section 4 concludes with a brief discussion.

## Materials and Methods

Here we introduce notation and motivate the methodology through a simple simulation study. Consider a set of *N* = 30*K* nucleotide sequences *X*_*n*_ where each sequence is of length *I*_*n*_ = 200 and composed of bases *A, C, G, T* drawn from some genome background probabilities (e.g. [.3,.2,.2,.3]). Randomly label half of the *N* sequences *Y*_*n*_ = 1 and the remaining *Y*_*n*_ = 0. For 98% of the positively-labeled cases, insert the sequence *GGGGGGG* at position *i* ∈ 1 : *I*_*n*_ with *i* drawn uniformly at random. Conversely, insert the sequence *CCCCCCC* into the negatively-labeled cases with a similar uniform-location probability. We wish to train a binary classifier to predict the associated label *Y*_*n*_ ∈ {0, 1} for a given sequence *X*_*n*_. Of course, under this framework, perfect model accuracy would be obtained if an oracle could encode a binary feature denoting the presence/absence of *GGGGGGG* in each sequence (or similarly, the *CCCCCCC*). The *discovery* and *representation* of such a sequence, however, is what interests us. In other words, can our model directly learn the predictive subsequences without defining features *a priori* or requiring post-hoc interpretation procedures?

We utilize techniques from the deep learning literature to form such a feature finder. Specifically, we consider the convolution operator employed in convolutional deep neural network (CNN) architectures [14]. Consider a set of *J* convolutional filters where the *j*^th^ convolutional operator computes the inner product between a weight matrix Ω^*j*^ and an observation matrix *X*_*n*_ at each position sliding along user-specified dimensions. These convolutional filters, or patches, are particularly suited for learning local spatial relationships and when employed in deep learning are stacked together, potentially in the hundreds within a single layer, from which the activations produced by each convolutional filter are fed as input into subsequent deeper layers. In genomics applications each first-layer filter Ω^*j*^ is a 4 × *L*_*j*_ matrix of weights 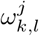 for *k* ∈ {*A, C, G, T*} and *l* ∈ 1 : *L*_*j*_ convolved upon a one-hot encoding of the input sequence *X*_*n*_. That is, each *X*_*n*_ is transformed from a nucleotide string of length *I*_*n*_ into a binary matrix of size 4 × *I*_*n*_ with rows corresponding to each base and column *i* denoting the presence/absence of a nucleotide at position *i* ∈ 1 : *I*_*n*_. Generally some sort of pooling operation is performed such that either the maximum (max-pooling) or average (average pooling) is selected within a small window, effectively reducing the parameter space and alleviating observational noise. We may write the model explicitly under the logistic link function with max-pooling performed over the entire input sequence as 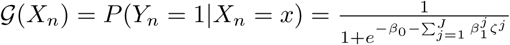 where 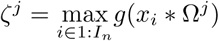 indicates a max-pooled convolution operation. The convolutional operation itself is explicitly defined as 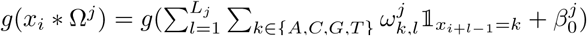 with *g*(*·*) representing the sigmoidal activation function in our experiments. We note it is these Ω^*j*^ matrices which contain the weights which collectively capture the sequence motifs and are often visualized as sequence logos through the methods described in the introduction.

For this first simulation study, we arbitrarily set *J* = 4 and *L*_*j*_ = 12 for all *j*. We restrict our model to simply the maximum value of the convolutional operator per filter as this represents the best match, or similarity score, between the motif and the sequence, and is also readily interpretable while maintaining parsimony. The four activation values produced from the four filters may be thought of as constituting the input features, or design matrix, to a logistic regression formulation. Any additional predictors may of course be included as well. It should be noted that all subsequent formulations may be extended to any number of layers as the model described is equivalently a single convolutional layer and single connected layer CNN (i.e. a *shallow* CNN). This vanilla CNN provides the baseline comparison and is depicted in panel A of Fig. 1. Of particular importance is to note that the weights, 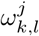, (middle row of sequence logos within panel A) are unconstrained. Thus we also provide the sequence-logo motifs calculated as described in the introduction in the bottom row of panel A with the threshold set at 0.75 × max(activation) per filter. The background nucleotide probabilities used when calculating, displaying, or learning any PWMS/IGMs presented herein are taken to be uniform (i.e. *b*_*k*_ =.25, *k* ∈ {*A, C, G, T*}). While our method and software implementation allow for non-uniform probabilities, motifs such as those downloaded from JASPAR [10] are generally calculated and visualized against a uniform background. We opt to follow suit and note that learning motifs as IGMs against a uniform background is the simplest and quickest option in our implementation. Plots of the weights are also readily interpretable as sequence logos. Details are provided in the supplementary materials.

The top row box plots depict the test set activation differences for each filter broken down by true label (*Y* ∈ {0, 1}, left versus right, respectively), as well as the associated 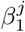 coefficients in red. These 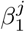 may be interpreted as effect size estimates for each motif. Not striking is the observation that the sign of the 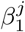 coefficients associated with filters *j* = 1, 2, 3 is negative while it is positive for filter 4. The sequence logos indicate that, as expected, the strings of cytosine nucleotides are highly predictive for negative sequences while the string of guanine nucleotides is highly predictive for positive sequences.

Our first contribution is illustrated in panel B of Fig. 1: we constrain the model weights during the training procedure to encourage motif interpretability. Specifically, we restrict the individual filter weights 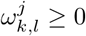 and their associated offsets 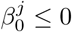, and additionally, re-scale the weights column-wise to maintain a valid information-theoretic relationship peri-training. The constraint on the offset weights 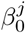 for each filter to be strictly non-positive is incorporated to improve the interpretation of the filter activations: consider that the minimum activation value, 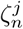 for observation *n* and filter *j*, attainable without a bias offset 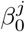 and under the sigmoidal activation would be 1*/*2. Quite simply the addition of a negative offset allows the value of 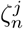 to decrease to zero. The middle row of Fig. 1B highlights the utility of the weight constraints by plotting the weights directly; no input test set observations or post-hoc optimization procedures were required. The filters maintain the strong class-discrimination as evident in the top row box plots, however there appears significant redundancy as the same 10-mer motifs were learned by two filters each. Thus our second contribution, also aimed at encouraging model interpretability, is to regularize the weights during the training procedure. We utilize the sparse group lasso penalty with each filter defined as a group [24]. L1 regularization (i.e. the so-called *lasso* penalty [25]) on the filter weights pushes non-informative weights to zero and may be interpreted as encouraging a KL divergence of 0 between the observed distribution and the background probabilities. We consider the sum of all weights in a filter as a measure of total motif information gain and these are the values regularized via the group lasso penalty which enourages irrelevent motifs to contain zero information gain and discourages correlated motifs. We detail these regularization schemes and interpretations in the supplementary materials.

Panel C of Fig. 1 shows the results of such a regularization scheme applied to the vanilla CNN in panel A. As desired, two of the filters now contain zero information gain and their associated effect estimates 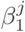 and offsets 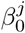 (not pictured) are also zero. These filters may be discarded without impacting model performance. Finally, panel D shows the results of utilizing both the constraints and the regularization scheme. We see the weights perfectly recapitulate the inserted 8-mer sequences and in fact illustrate the motif more clearly than the approach based on test set observations (bottom row). We note that all models achieved near-perfect predictive accuracy (98%) and were trained for five epochs. Parameter tuning was performed to identify suitable values for the regularization penalties. While we only present the results for learning IGMs, learning PWMs is also possible and included in our software implementation. The reader is encouraged to view the supplement for a brief discussion on the implications of learning IGMs or PWMs.

## Results

### Simulation Study

The toy example previously described is useful for illustration but it represents an unrealistic situation in which nearly all observations contain an optimal representative motif. We therefore consider a second and more challenging simulation study and base the methodology on that laid out in [22]. Specifically, we utilize the simdna package [11] to generate 100K nucleotide sequences of length 200bp sampling from motifs with less degenerate distributions. We sampled from three motifs: MYC, CTCF, and IRF^1^, where positively-labelled sequences (*Y* = 1) contain 0-3 occurrences of each motif. 100K negatively-labelled sequences (*Y* = 0) were generated from random genome background with 40% GC-content. Additionally, 10% of the observations were shuffled between positive and negative classes to increase the difficulty of the learning procedure. The top row of Fig. 2B shows the target sequence logos for the three sampled motifs (*MYC known1, CTCF known1, IRF known1*) embedded in the positive cases. These motifs are the subsequences we wish to learn.

**Fig 2.**
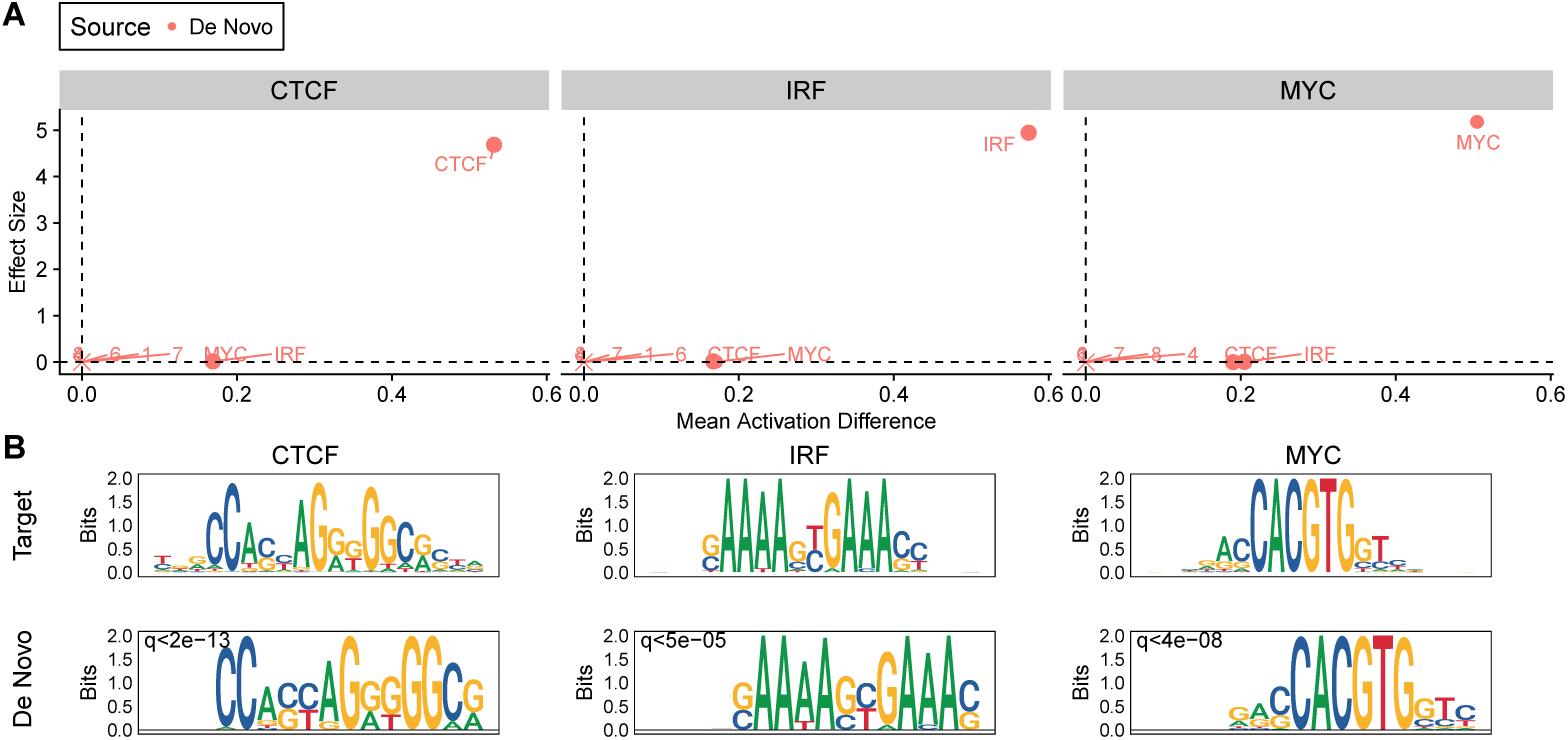
*De novo* motif simulation results. Positive cases contain 0-3 insertions of target motifs: MYC, CTCF, IRF (top row panel B) whereas negative cases are sampled from random background. All first-layer convolutional filter weights are initialized randomly and motifs are learned *de novo*. **A.** Filter motif effect size (Y-axis) against mean activation difference between class labels (X-axis). × indicates zero information gain motifs and the associated numeric label indicates the filter number (see Supplementary Fig. S1. **B.** Target sequence-logo motifs (top row), Learned *de novo* motifs (bottom row). Q-values reported from running Tomtom motif comparison tool. [8].

#### Multi-dimensional output model

A regularized and constrained CNN (as described in the previous section) was trained via stochastic gradient descent for twenty epochs with the learning rate initially set at 0.02. *J* = 8 first-layer convolutional filters were randomly initialized following a uniform distribution on the interval (0, 0.5). Logistic loss plus the regularization terms was minimized over the three motif classes. Thus the target output vector *Y*_*n*_ for observation *X*_*n*_ is a 1 × 3 binary array with each entry indicating the presence/absence of motif *j*. Sequences containing no motif instances (i.e. purely random background) are labelled *Y*_*n*_ = [0 0 0] whereas a sequence containing, for example, the motifs MYC and CTCF but not IRF would be labelled *Y*_*n*_ = [1 1 0]. Under such a formulation, 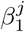 is no longer a 1 × *J* vector but a 3 ∈ *J* matrix with rows corresponding to entries in the target array *Y.* Fig. 2 panel A plots the fitted 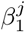 estimates (Y-axis) against the mean activation difference between test set observations containing *any* motif occurrences (i.e. any entry in the 1 × 3 binary array is greater than zero) and test set observations containing *no* motif occurrences (all entries exactly zero). Faceting corresponds to rows in the 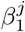 matrix such that the heading CTCF represents the CTCF target class, the IRF heading represents the IRF target class, etc. It is indeed reassuring that a single filter exhibits both the largest mean activation difference and the largest effect size within a facet, and whence visualized as a sequence logo (panel B bottom row) this filter recapitulates the desired target motif (panel B top row). Five of the eight filters have associated *β*^*j*^ coefficients equal to zero across all facets as well as zero activation difference. These filters may be removed from the model without impacting predictive accuracy and, as evident in Supplementary Fig. S1, are zero information gain motifs, thus indicating the effectiveness of the regularization. Only the three filters with non-zero information gain, effect size, and mean activation difference need to be retained in the model. The weights composing these filters are shown in panel B of Fig. 2 with associated Q-values from running the Tomtom motif matching tool [8]. In all three cases the most significant Q-value is the desired target motif (Supplementary Fig. S2). We label the points in panel A with the most significant Tomtom match and note that due to the construction of the simulation (sequences may contain two or even three different motifs), the mean activation difference is non-zero for two motifs in each facet however the effect size estimate is zero. Thus the two off-target *de novo* learned motifs are uninformative for prediction of a given target class, however approximately one-third of sequences may of course contain either (or both) of the motifs.

#### Single-dimensional output model

While often considered in the literature the multiple-output model previously described is of little use in practice; rarely would such labels exist denoting the presence/absence of each motif. Indeed it is these motifs which we wish to learn and thus a single-output model is of more practical use. Such a model formulation might arise from, for example, a ChIP-seq experiment in which sequences extended from called peaks would be labelled as *Y* = 1 while sequences of equivalent length would be drawn from random genome background and labelled as *Y* = 0. The analytic goal of such a formulation would be, again, discovering which motifs (perhaps even beyond those ChIP-ed for) are abundant in the peaks versus the background. For this reason we collapse the 1 × 3 target vector into a single value denoting the presence of *any* motif (*Y*_*n*_ = 1) versus the absence of *all* motifs (*Y*_*n*_ = 0) and highlight a use case for our method. We show how one might initialize filters based on annotated motifs from a database and also learn any extra motifs *de novo*.

We consider two models to highlight this use case: Model 1 initializes filters based on the 579 previously-annotated motifs found in the 2018 CORE vertebrates non-redundant JASPAR database and holds these filter weights fixed throughout training [10]. Such a use case might arise when one does not have *a priori* knowledge of which motifs may be present in the sequences and wishes to estimate the prevalence of previously-annotated motifs. Model 2, on the other hand, initializes two filters with the JASPAR MYC and CTCF motifs and tackles the issue of discovering a motif which is present in the data but not the motif database (in this case, the IRF/IRF2 motif). To achieve this we simply remove the IRF2 motif from the filter initialization and try to learn it *de novo*. We initialize two filters uniformly at random on the interval (0,.5) and learn the motif directly. Such a use case might arise during a specific TF ChIP-seq experiment when one believes several previously-annotated motifs may be present but also wishes to learn unannotated motifs *de novo*. In the first model we impose sparse regularization via the L1-norm on the 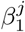 coefficients to encourage those motifs with little-to-no abundance to exhibit an exactly zero effect size 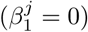 while in the second model we impose the regularization strategy outlined in the previous section to discourage redundancy.

Supplementary Fig. S3 panel A plots the estimated effect size 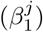 against the mean activation difference between classes in the test set for each JASPAR-initialized convolutional filter (post-sigmoid transformation). It is evident that, of 579 JASPAR-initialized filters, only four exhibit an effect size greater than zero and three of these are the true motifs used in the data simulation (CTCF, IRF/IRF2, MYC). Panel B shows that the fourth, MAX::MYC, is nearly identical to MYC and one may include only a single instance of these during the initialization procedure. We conclude that 575 of the 579 convolutional filters (motifs) may be discarded with no effect on model performance. Similarly, Fig. 3 showcases the results for Model 2, in which the IRF motif has been removed. Of note is the large activation difference and estimated effect size for the *de novo* learned IRF motif (Panel A). As desired, the estimated effect size is of an equivalent magnitude as the JASPAR-initialized motifs.

**Fig 3.**
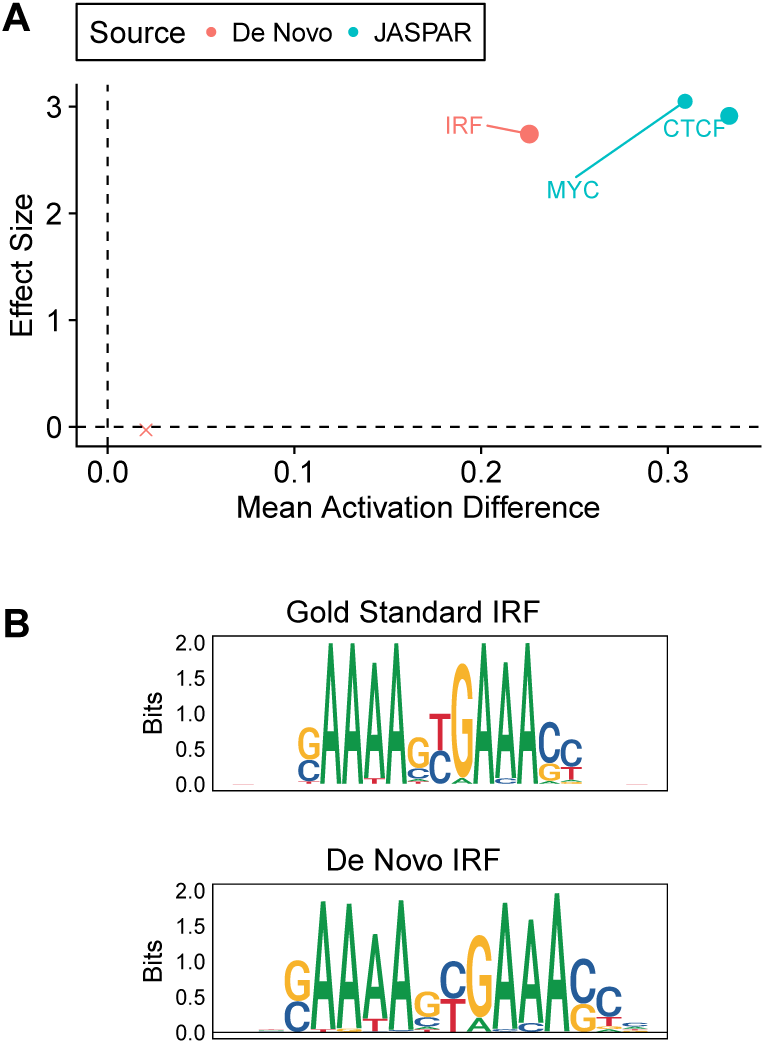
Simulation model 2. The two first-layer filters initialized to JASPAR-annotated CTCF and MYC motifs, however the IRF motif must be learned *de novo*. **A.** Filter motif effect size against mean activation difference between class labels by motif source indicates equivalent magnitudes of effect size for both the JASPAR filters and the *de novo* filters. The red cross indicates a low-information gain *de novo* filter that may discarded without affecting model performance. **B.** Gold standard IRF motif (top) exhibits high similarity with *de novo* IRF motif (bottom).

Comparing the gold standard embedded motif (left-hand sequence logo of panel B) with the *de novo* IRF motif it is evident how similar these motifs are and how one might utilize our tool for learning motifs *de novo*. We note that, in addition to simply initializing and fixing filters with JASPAR-annotated motifs, one need not hold these fixed during training and may instead choose to update the individual filter weights (motif position-probabilities) to both improve model fit and compare the updated motif with the original motif. We leave this for future work.

#### Latent variable interpretation

Under the sigmoidal activation function applied to the convolution output (activations) from Model 2, each 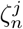 is interpretable as the probability that sequence *n* contains motif *j*, i.e. 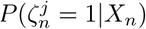. We sought to assess the accuracy of this latent variable interpretation in Fig. 4A-C on a held out test set in which we did not randomly shuffle 10% of the observations between classes. We find the 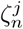 representations are extremely accurate, achieving ≥ 98% accuracy and an area-under-the-precision-recall curve of ≥ 0.99 for all three motifs (MYC, CTCF, IRF).

Pooling information across all motifs slightly diminishes performance, especially for the sequences containing only a single unique motif (combined-model accuracy for sequences containing a single motif ranges from 0.617 – 0.816 yet accuracy is perfect for all sequences containing two or more unique motifs). One should take care to note the imbalance in the individual motif comparisons relative to the balanced dataset (1:4 versus 1:1 positive:negative cases, respectively). Additionally, in panel D, we explored the possibility of using Monte Carlo Bernoulli draws with probability based on the individual 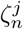, and using the realizations to fit a logistic GLM. We find all coefficients are statistically significant at the *α* =.05 level across all simulations (*m* = 1000), however the fitted effect sizes underestimate the true effect as fixed by the simulation, presumably due to the collinearity of the predictors. Panel E plots two dimensions from multi-dimensional scaling (MDS) performed on the activation values from the retained filters (three in total), highlighting a separation between both the class labels (*Y* = 0*/*1) and, to a lesser degree, the separation within the positive cases due to the underlying embedded motifs.

**Fig 4.**
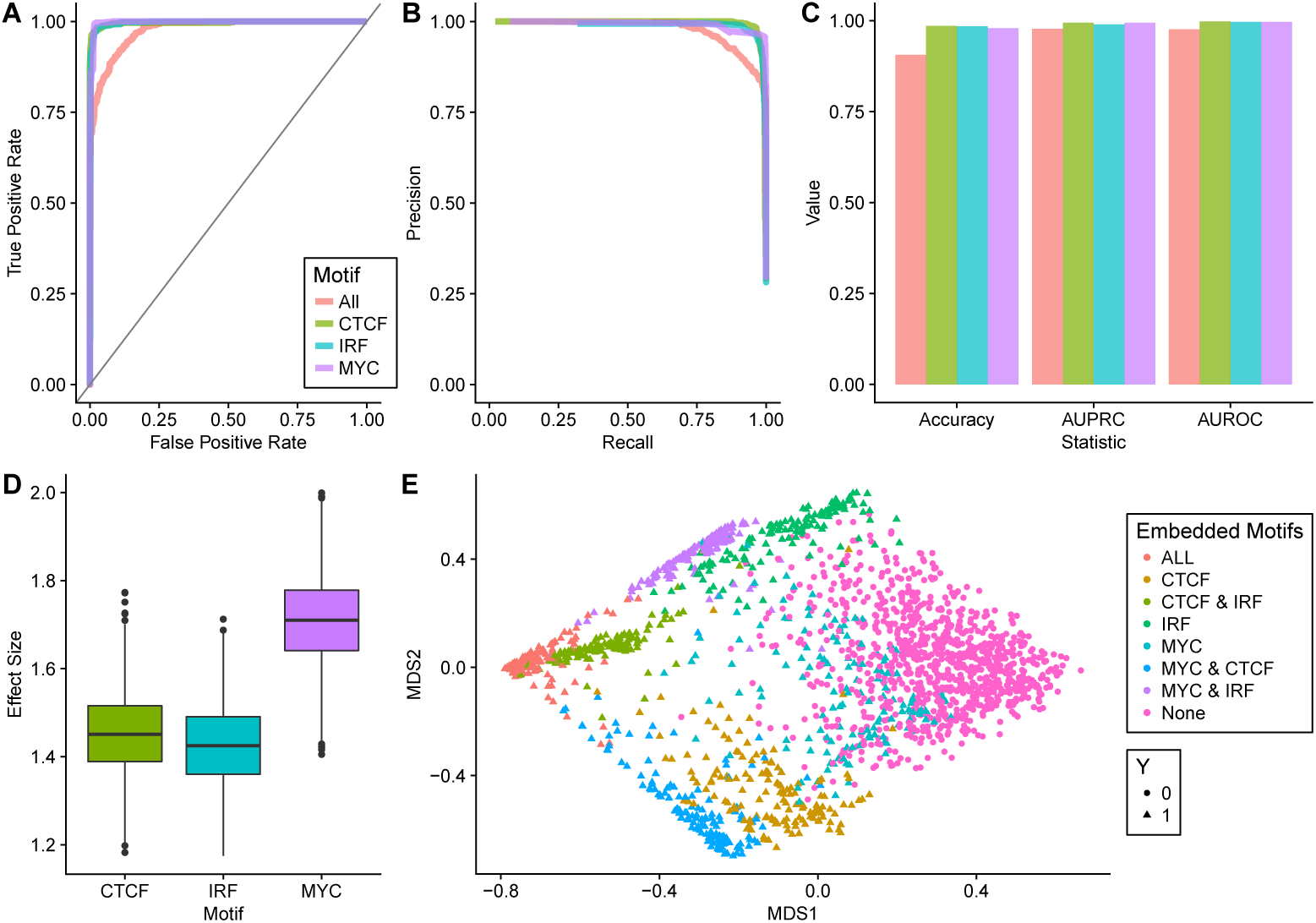
Model-based evaluation of simulated motif presence. **A-C.** ROC curve, precision-recall curve, and prediction statistics evaluating correctness of calling individual motifs present within test set sequences based on max-pooled filter activation values. The *All* label simply combines the three filters in a logistic regression model to evaluate presence of *any* motif versus absence of *all* motifs. Dashed lines represent using the true, sampled-from motif whereas solid lines represent the *de novo*-learned motifs. **D.** Effect size estimates for each motif based on Monte Carlo realizations treating the max-pooled filter activation values as Bernoulli random variables. **E.** Multi-dimensional scaling using the max-pooled filter activation values as features.

### Data Application

We applied our method to *in vivo* transcription factoring binding data from the ENCODE Consortium [6]. Specifically, following the protocol described in [23] with slight modifications, we downloaded CTCF TF ChIP-seq data for the Gm12878 cell line. 100bp windows were extended from the called peaks for the positive cases while negative cases were obtained by removing from all DNase peaks in Gm12878 the top 150K relaxed TF peaks called by SPP at a 90% FDR [13]. Peaks originating from chromosome 1 were utilized for the training set and peaks originating from chromosome 2 were utilized for the validation set. We down-sampled the negative cases (genome background) during the training procedure however did not down-sample cases in the validation set.

Fig. 5A showcases the utility of our approach: we begin by initializing the filters with all possible JASPAR motifs. We denote this as the shotgun approach as many of the motifs miss the mark (e.g. left-hand panel of A, the vast majority of motifs exhibit both 0 mean activation difference and 0 effect size). We discard all filters from the model which do not have both an estimated effect size greater than 0.01. In the case of this analysis, five motifs were retained and indeed it is reassuring to see both the CTCF motif and the CTCF reverse complement (CTCF RC) as retained. We train this model for 10 epochs to obtain fitted values for all filters’ 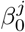 and 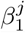, and then initialize a *de novo* model with these five filters (and their associated *β*) fixed, but also eight filters randomly initialized. These latter filters will be used to learn the *de novo* motifs. We train this model for 30 epochs, this time utilizing both the regularization tactics and the weight constraints, and report the estimated effect size and mean activation difference for the *de novo* motifs in the middle panel. We calculate the information gain of each motif as the sum of all weights in the filter and provide this value as the size of the associated point. We find six filters to contain zero information gain and thus we discard these filters. Our final model then makes use of the five JASPAR filters and the two *de novo* filters, and we train this model for another 30 epochs to obtain values for each 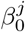 and 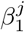, as well as the overall offset *β*_0_. The right-hand panel of Fig. 5A illustrates both the high information gain of the *de novo* motifs, as well as the larger effect size and mean activation difference. Visualizing all the motifs in Fig. 5B sheds light on what the *de novo* filters have learned: namely slightly altered representations of the CTCF and CTCF RC motifs. In fact, we find the effect of the leading G to be amplified in the *de novo* 2 motif relative to the CTCF known motif, and, correspondingly, the trailing C in the RC to be amplified. This suggests the subsequence GCGC is more abundant than expected by the CTCF JASPAR motifs. Similarly we note the deletion of a rather uninformative position in the motif (position 18 in CTCF RC and position 2 in CTCF). We finally provide test set accuracy statistics and illustrate via MDS the class-separability of sequences using these seven activation values as features.

**Fig 5.**
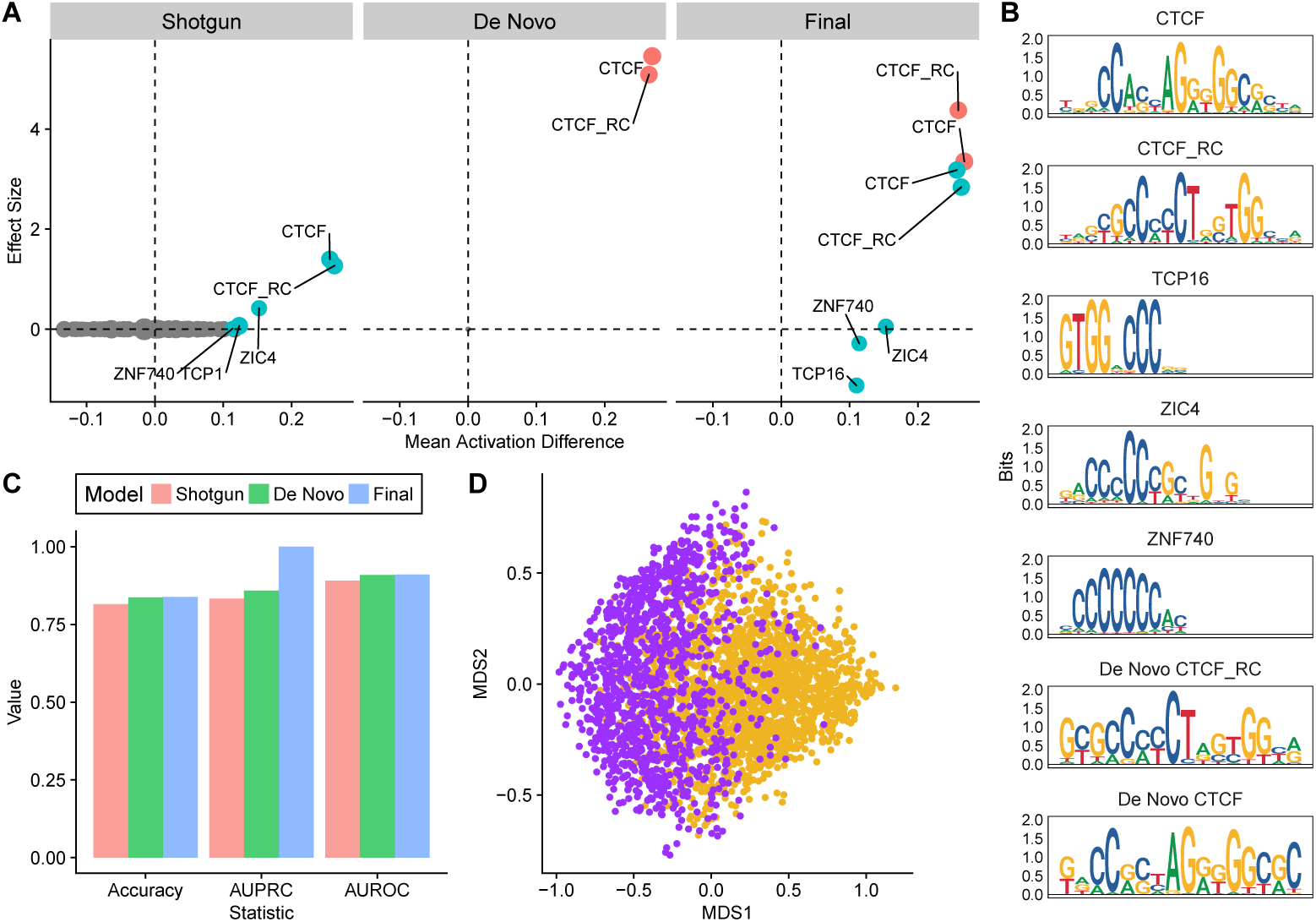
ENCODE pipeline results. **A.** Motif effect size against mean activation difference between class labels. The *Pipeline* model includes all 2018 JASPAR-annotated motifs (core vertebrates, non-redundant). Any motifs with associated effect size *<.*01 are discarded and subsequently held fixed while eight *de novo* motifs are learned (center panel). The *Final* model discards uninformative *de novo* motifs and refits effect size estimates with filter weights fixed. **B.** Collection of motifs selected from *Final* model plotted as sequence-logos. **C.** Prediction statistics evaluating classification based on model. **D.** Multi-dimensional scaling using the max-pooled filter activation values from the *Final* as features, colored by label (purple: *Y* = 1, yellow: *Y* = 0).

## Discussion

Our proposed model directly learns sequence motifs such that interpretation of the convolutional filters is not reliant upon test set observations and the weights comprising the filter are directly interpretable as information gain measures or position weight (log-odds) measures. We address the issue of filter redundancy along with interpretability by incorporating weight constraints and regularization techniques. The weight constraints limit the range of the individual filter weights to restrict their values to be directly interpretable as IGMs/PWMs while the regularization scheme encourages learning non-redundant and high-relevance motifs.

To the authors knowledge this is the first method capable of incorporating previously-annotated sequence motifs in a CNN framework, similar to [19] but with the ability to learn motifs *de novo*. Notably the method achieves this by leveraging IGMs/PWMs as convolutional filters and ensuring the *de novo* motifs are valid information gain/position weight measures. Interestingly, other motif measurement systems such as the position probability matrix (PPM) may also be used as convolutional filters although several changes must be made. First, the regularization scheme must be adjusted. In the case of the PPM, filter weights would need to be regularized around their expected background frequencies (e.g. 0.25). Further, all weights would require a column-wise sum to unity, thus all weights would need to be initialized under such a condition and enforced throughout the training procedure. Additionally, any low-information gain motifs would be those filters with all weights centered at their background frequencies, and thus summing all weights within the motif would not constitute a measure of information gain since all sums would be the same, regardless of the amount of information gain contained. One would likely transform the PPM into an IGM in order to quantify importance as measured by the KL divergence. The same holds for using the PWM, as the negative values associated with the low abundance nucleotide positions would overwhelm the sum calculation.

Like any regularization method, parameter tuning is essential and as the number of parameters to tune increases, so does the difficulty in finding suitable values. This issue, however, is ubiquitous with deep learning techniques and does not affect our method any more than usual. Furthermore, as our primary concern is more with learning discriminatory features and less with predictive accuracy, we find parameter tuning to act as a sort of interpretability sieve; under stricter regularization only the most discriminatory features will appear at the cost of predictive accuracy while under laxer regularization predictive accuracy may improve at the cost of lesser filter interpretability. Indeed this trade-off epitomizes the divide between traditional statistical techniques and machine learning methods, however once discriminatory motifs/features have been learned one may refit a more complicated (i.e. deeper) model to attain improved predictive accuracy. One may even desire to learn motifs *de novo* as IGMs, and then refit the model using PWMs given the one-to-one correspondence between the two.

The proposed methods may be useful for several interesting genomics applications; namely any application requiring the need to learn differential sequence motifs. Examples include DNase-seq and ATAC-seq footprinting. We leverage deep learning infrastructures to provide a more succinct set of features and abandon the traditional machine learning paradigm stating that higher accuracy is paramount. We instead focus on a simple modelling framework which provides end-to-end interpretation throughout. Our methods are implemented in an R package available at https://github.com/mPloenzke/learnMotifs which relies upon Keras [5] for all of the heavy lifting.

## Acknowledgments

We wish to thank Chinmay Shukla, Mingxiang Teng, Patrick Kimes, Keegan Korthauer, and Alejandro Reyes for their valuable feedback during the preparation of this manuscript.

## Supplementary Methods

### From nucleotide sequences to information gain

A position frequency matrix (PFM) tabulates the frequency of each nucleotide at each position in a set of reads. For example, consider 100 sequences each of length six letters composed of the nucleotides {*A, C, G, T*}. A single example sequence could be *ACCTAG*. Under a uniform distribution, one would expect 25 of each nucleotide at each position and the resulting PFM would be:

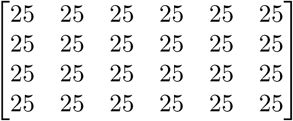

The top-left entry corresponds to the count of *A* nucleotides in the first position. Dividing by the column sums yields a position probability matrix (PPM), of which each column defines a multinomially-distributed random variable. Denote this r.v. *P*_*c*_ for column *c* and for the example consider the PFM and resulting PPM below:

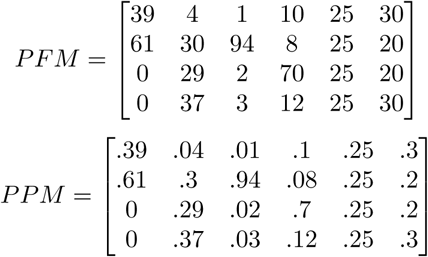

If one observed counts as tallied in column three of the PFM, then one might expect to reject a frequentist null hypothesis of uniform nucleotide probability (*P*_3_ ∼ multinomial(1,.25,.25,.25,.25)). A likelihood ratio test could be used to perform such inference with the log-likelihood of observing the data under the null taking the form [4]:

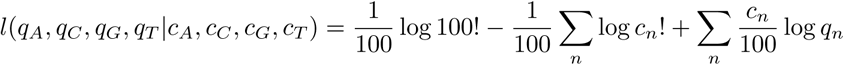

where *q*_*n*_ denotes the null probabilities (0.25) for nucleotide *n* and *c*_*n*_ denotes the observed count (in the PFM). Following the derivation provided by [21] and making use Stirling’s approximation for large *N* (log *N* ! ≈ *N* log *N - N*) gives us:

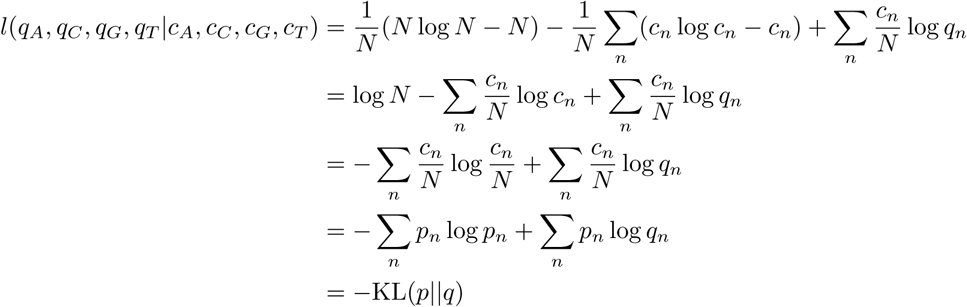

Here *N* denotes the total number of sequences observed (100 in the example) and *p*_*n*_ denotes the observed PPM values for nucleotide *n* ∈ {*A, C, G, T*} calculated from the PFM. The last line follows by the definition of Kullback–Leibler (KL) divergence, a measure which has many interpretations including the negative observed log-likelihood, the relative entropy or information content, and in the bayesian paradigm the information gain from using the posterior probabilities *p*_*n*_ relative to *q*_*n*_. Rearranging the last line reveals how the this quantity takes into account the background distribution:

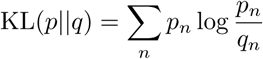

This is a weighted sum of the log-odds 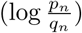, terms which define the position weight matrix (PWM). Under a uniform background, the equation above simplifies to 2–∑_*n*_ *p*_*n*_ log *p*_*n*_ and in our example gives the following PWM:

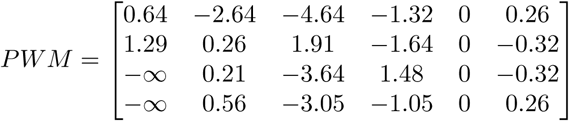

Each entry is upper bounded by 2 due to the choice of the uniform background but the entries are not lower bounded. The absolute magnitudes of the column sums indicate how different the probabilities are and are also unbounded. Pseudo-counts may be added to the PFM or PPM to remedy the -∞ terms. When done so by adding 0.5 to the zero entries in the PFM results in the following PWM and KL divergence:

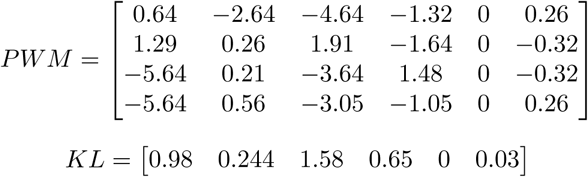

As expected the KL divergence is zero in the fifth column, small in the sixth column, and largest in the third and first columns. In the bayesian paradigm, the information gained by using the posterior probabilities depicted in the PPM relative to a uniform prior would be largest in the third column and zero in the fifth. One may lastly obtain the weights shown in the sequence logo plots [20] by multiplying each value in the PPM by its corresponding column in the KL divergence vector. We term this matrix the information gain matrix (IGM). Many other names would suffice including the sequence logo matrix.

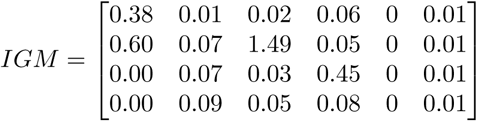

Of course the IGM and PWM contain the same information and are simply rescalings; one may start with a PWM and calculate an IGM and vice versa. Indeed one may transform a PWM/IGM based on background probabilities *q*_*n*_ into a PWM/IGM based on background probabilities *r*_*n*_ simply by backing out the *p*_*n*_ and recalculating the quantity of interest.

### Model formulation

We develop our model based on the convolution operator employed in convolutional deep neural network (CNN) architectures [14] considering the PWMs and IGMs as the filters. Since each filter is itself directly interpretable as a sequence motif, we may initialize (or fix) the filter values with those of previously-annotated database motifs directly. We may also attempt to learn the motifs *de novo* given the weights maintain an equivalent interpretation.

Consider a set of *J* convolutional filters where the *j*^th^ convolution operator computes the inner product between a weight matrix Ω^*j*^ and an observation matrix *X*_*n*_ at each position sliding along user-specified dimensions. These convolutional filters, or patches, are particularly suited for learning local spatial relationships and when employed in deep learning are often stacked together in the hundreds within a single layer, from which the activations produced by each convolutional filter at each position are fed as input into subsequent deeper layers. The choice to use these filters to learn sequence motifs is motivated by the work of [2,9,29]. In these genomics applications each filter Ω^*j*^ is a 4 × *L*_*j*_ matrix of weights 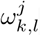 for *k* ∈ {*A, C, G, T*} and *l* ∈ 1 : *L*_*j*_ convolved upon a one-hot encoding of the input sequence *X*_*n*_. That is, each *X*_*n*_ is transformed from a nucleotide string of length *I*_*n*_ into a binary matrix of size 4 ×*I*_*n*_ with rows corresponding to each base and column *i* denoting the presence/absence of a nucleotide at position *i* ∈1 : *I*_*n*_.

We write the model explicitly under the logistic link function (i.e. the sigmoidal activation function), due to binary classification objective as follows:

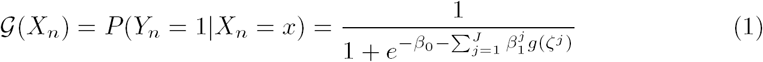

where *ζ*^*j*^ indicates a max-pooled convolution transformation for filter *j* at location *i*:

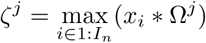

and *g*(*·*) is some activation function (e.g. linear). The convolution operation is explicitly defined for observation *X*_*n*_ = *x* at position *i* as:

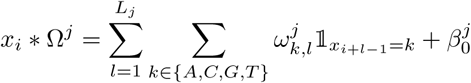

We again note that it is these Ω^*j*^ matrices which contain the weights which collectively capture the sequence motifs. We introduce this notation to motivate the merger of the convolutional filter as a linear predictor: consider the case when *J* = 1 and the weights are all fixed such that 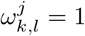 for *k* = *C, l* ∈ 1 : *L*_*j*_, and 0 else. This trivial filter does nothing more than compute the count of *C* nucleotides within a *L*_*j*_-bp sliding window for each observation and assigns the maximum sigmoidal-transformed value as a feature to be fed into a logistic regression model. Interestingly it does this by computing the similarity (cross-correlation) between each test sequence with the all-*C* motif. There is nothing ‘deep’ about this model and there are only two parameters to fit, *β*_0_ and *β*_1_, either via an iterative maximum-likelihood approach [17] or gradient descent. Moreover, the interpretations of these values directly correspond to the standard statistical interpretation of regression coefficients.

We now introduce the random variable 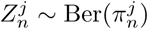 to denote the presence 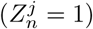 of motif *j* in sequence *X*_*n*_. 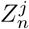 is unobserved and we wish to estimate it via:

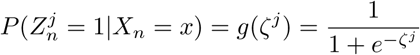

where *ζ*^*j*^ denotes the max-pooled convolution operator from above. Note that 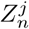 is computed simply by transforming the max-pooled convolution operation to be in [0, 1] via the sigmoid function. Recalling that the outcome variable *Y*_*n*_ is itself a random Bernoulli variable with probability *p*_*n*_ we may condition upon the hidden variables 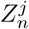 to rewrite equation 1 as:

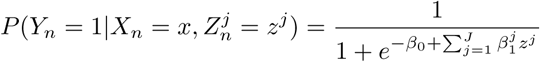

Thus *Y*_*n*_ is simply modeled via a transformation of a linear combination of hidden variables denoting the presence or absence of a given motif based on its maximum subsequence similarity. Of course there is no reason beyond parsimony and interpretability for the need to reduce the *j*^th^ feature map to a single maximum value (versus, for example, the sum of all elements, or the average within the first half of the sequence and a separate term for the average in the second half). We also note that one may encode any subsequent values from the *j*^th^ filter as further hidden states sharing the same motif filter to assess the additive impact of motif occurrences in this nested modeling framework. Finally, we note that one may consider a different distribution for the 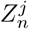, such as a Poisson to model to the count of a given motif or a linear link to model intensity, however we leave this for future work. We fit all model parameters via stochastic gradient descent in Keras [5] with a batch size of 64.

### Encouraging interpretability

Upon successful model training there are three interpretable quantities of interest in our model: 1) The set of filters Ω^*j*^ representing motifs, 2) their associated 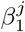 model coefficients of estimated effect sizes, and 3) the hidden variables 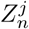 denoting the presence/absence of motif *j* in sequence *n*. We present this simple model in the text to encourage the interpretability of each model layer within the CNN terminology. One may opt for a more complicated (deeper) model while still maintaining the interpretation of filters as motifs. We detail the weight constraints and peri-training transformations below, along with some practical considerations for implementation.

### IGMs as filters

When representing the motifs (convolutional filters) as information gain matrices we restrict the individual filter weights 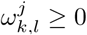 and their associated offsets 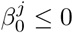. Additionally, the weights for a given filter at a given position must be valid information gain measures. To achieve this we restrict the column-wise sum to be less than or equal to 2 under uniform background and rescale the weights column-wise to maintain a valid information-theoretic relationship during training. This latter step is accomplished by rescaling the weights from information gain to probabilities by dividing each weight by its column-wise sum and subsequently converting back to information gain by multiplying each weight in the PPM by the column-wise sum of the expected self-information gain. A psuedo-count of 0.05 is added to entries whose column sum is less than 0.1 when converting to the PPM to control for cases in which a single, small weight in the column is non-zero and thus occupies a position-probability of 1. We perform this rescaling at the end of each training epoch. The constraint on the offset weights 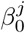 for each filter to be strictly non-positive is incorporated to improve the interpretation of the filter activations: consider that the minimum activation value, 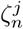 for observation *n* and filter *j*, could take without a bias offset 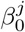 and under the sigmoidal activation would be 1*/*2. Quite simply the addition of a negative offset helps decrease the value of 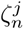 to 0 for certain *n*.

Under such a scheme we find that the learned filters may be interpreted as information gain and are directly comparable to previously-annotated database motifs in the form of IGMs. For this reason we consider the sum over the filter as a measure of information gain of the motif, which may equivalently be interpreted as the KL divergence. In addition, the associated 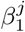 model coefficients are interpreted as the estimated effect size of the motif. Under the model described above, and within the context of the applications considered, the 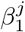 estimates translate to log-odds ratios. In experiments in which the negatively-labelled cases represent purely random genome background, we also constrain these 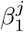 to be strictly non-negative. While the inclusion of such a constraint is surely debatable per the application at hand, when the task is to discover enriched motifs in the positively-labelled class, such as in the MYC-CTCF-IRF simulation, the constraint is justifiable as one does not expect to discover depleted motifs. Surely any motifs with negative effect sizes would, by construction of the simulation, be due to over-fitting or spurious learned features. For this reason we include the 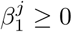 constraint in the data application, but do not in the first simulation presented with the *C*-motifs and *G*-motifs. In the latter case no such constraint is warranted.

### PWMs as filters

When representing the motifs as position wieght matrices an altered weight constraint scheme is used. We no longer require the non-negativity constraint on the 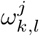 nor the 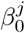 negativity constraint. We limit the upper bound of the individual weights to be less than or equal to 2 (again for the case of uniform background) and rescale the weights column-wise to maintain a valid distributional relationship during training. This latter step is accomplished by rescaling the weights from log-odds to probabilities by adding *log*_2_(*b*_*n*_) to each weight and raising two to this power. We subsequently convert the calculated probability back to a position weight by computing the log-odds. Again a psuedo-count of 0.05 is added to the zero entries to avoid values of negative infinity. We perform this rescaling at the end of each training epoch as in the IGM case. Under such a scheme the regularization is again interpretable, this time encouraging small and non-zero log-odds to shrink to zero. The filter-level regularization again discourages redundancy but this time does so on the log-odds scale instead of the information gain scale. Lastly, it is worth noting that backing the probabilities out during the weight rescaling performed for the PWM requires the addition operation and the power operation, while in the case of the IGM, the calculation simply requires the division by the column sum.

### Practical considerations

All sequence motifs have been represented as information gain matrices (IGMs) in this manuscript however this need not be the case and our software implementation provides support for initializing and/or learning motifs as PWM representations. As previously described, there is a one-to-one correspondence between the two measures some notes should be made on the implications between choosing one representation over the other. Notably the use of a PWM tends to spread out the activation values due to the negatively-unboundedness of the log-odds computed on multinomial probabilities. This leads to an asymmetry in how the sparsity in interpreted since extremely small position probabilities will never attain zero probability (*-*∞ position weight). The same is not true for the IGM representation as low position probabilities indeed attain zero. In other words, in the case of the IGM the sparsity encourages position probabilities to not only attain exactly 0.25 but to also attain smaller values. In the case of the PWM the sparsity exclusively encourages position probabilities of 0.25. Another consideration is that interpretation and weight visualization is a bit more nuanced when utilizing PWMs because the weights may be both positive and negative. Whereas added weight constraints were used in the case of the IGM to encourage interpretability, the same is not justifiable for the PWM. Additionally, in light of the large influence of increasingly-negative position weights, one may opt to abandon the latent variable interpretation presented herein and instead use a ReLU activation function to alleviate this. Undoubtedly it is a decision best determined by the problem at hand and the goals of the analysis.

A uniform background assumption simplifies the training procedure because the maximum value in the weight constraints is simply two across the board. Under a different background probabilities assumption, we do not include a maximum constraint while performing gradient descent, and instead recommend rescaling the weights after each batch (as opposed to epoch). This procedure is computationally more expensive and one may instead train all weights under the uniform background assumption and rescale them after the fact.

### Regularizing redundancy

Despite encouraging interpretable learning through weight constraints, the filters tend to learn correlated and often redundant features, with this being related to the hyper-parameter *J* determining the number of such motifs to learn *de novo*. We remedy this issue through regularization, specifically by incorporating a sparse group lasso penalty into the formulation to encourage learning highly predictive and distinct motifs [24]. Our regularization scheme may be expressed as:

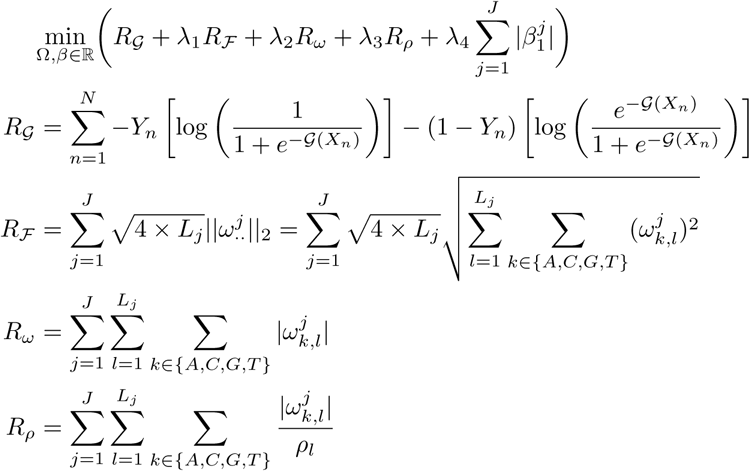

Model training proceeds by trying to find the weights which minimize this sum. In other words, gradient descent is performed to minimize this sum. *R*_*𝒢*_ simply denotes the standard logistic loss function and the *λ* parameters dictate the trade-off between minimizing this loss at the cost of each regularization penalty, respectively. *R*_*ℱ*_ encourages filter-level group sparsity with the L2/L1 norm [28], *R*_*ω*_ encourages nucleotide-level sparsity with the L1 norm, and *R*_*ρ*_ encourages motifs to form near the center of the filter with a location-specific penalty *ρ*_*l*_. Recall in the genomic data applications we consider each filter is a 4 × *L*_*j*_ matrix, with *L*_*j*_ assuming values around 8-16 depending on the specific problem. We wish to discourage learning shifted versions of the same motif and so we set the vector *ρ* to be a concatenation of a sequence of decreasing values beginning at *λ*_3_ and ending at 0 of length of equal to *L*_*j*_*/*2, concatted with a reversed version of the same sequence (i.e. increasing values from 0 to *λ*_3_). Thus sparsity is more strongly encouraged at the outer positions of the filter than towards the middle, in turn discouraging redundant and shifted versions of the same motif. Finally, we penalize the L1 norm of the 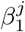 and include this regularization penalty in many of the models considered throughout the text. This penalty simply pushes effect size estimates to 0 and is often employed in CNNs. While group sparsity has surely been implemented in the context of image analysis (for example, [15,26,27], this is the first instance the authors are aware of using this regularization framework on genomic data. The regularization scheme in total pushes many individual weights to zero and discourages spurious motifs by pushing entire filters to zero except for those attaining suitably large KL divergence (or log-odds in the case of the PWM).

## Supplementary Figures

**Fig S1.**
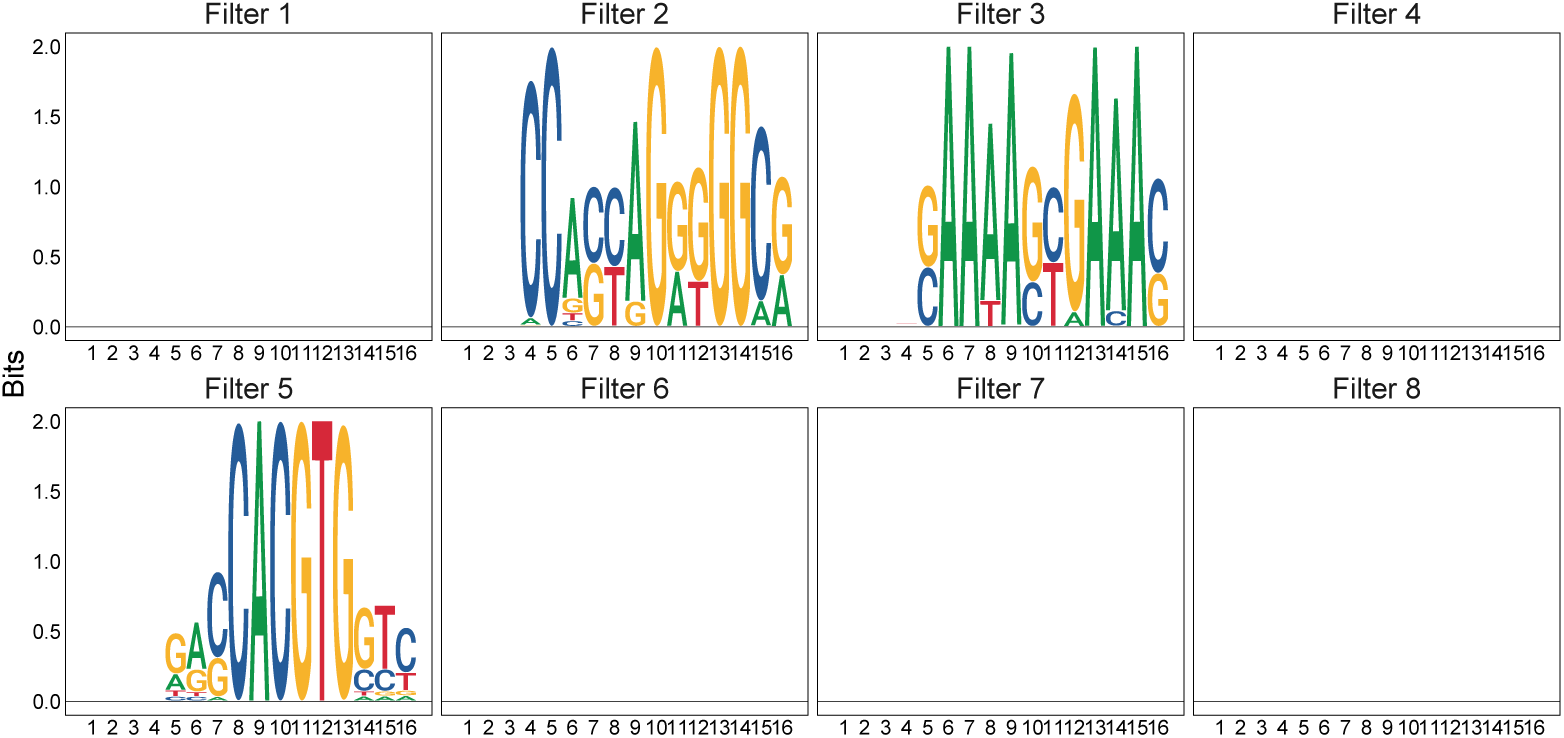
All *de novo* motif simulation filters. All first-layer convolutional filter weights trained on the CTCF-IRF-MYC simulation dataset. Many weights are regularized to zero and thus do not contribute to the model performance.

**Fig S2.**
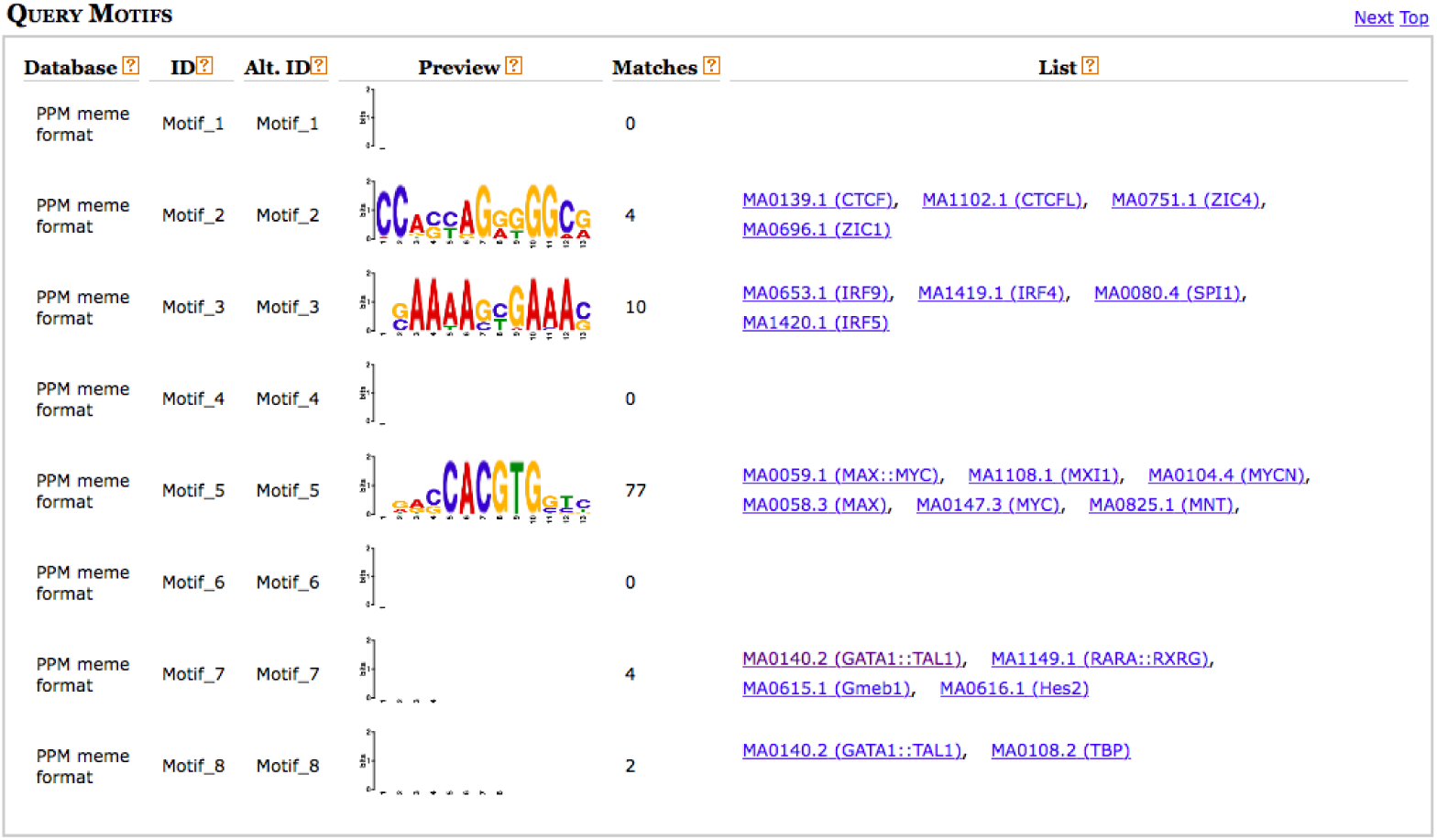
*de novo* motif simulation Tomtom query. Results from running the Tomtom motif similarity tools [8] querying learned motifs against the 2018 JASPAR Core Non-vertebrates database [10].

**Fig S3.**
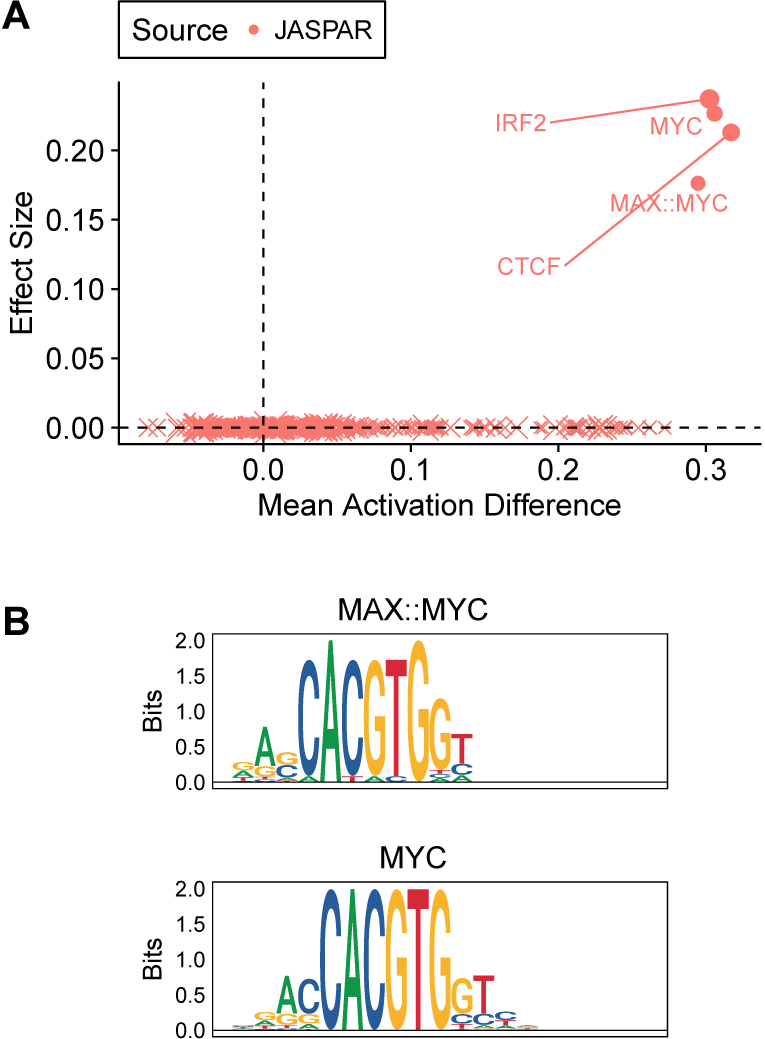
Simulation model 1. All first-layer convolutional filters initialized to 2018 JASPAR Core Non-vertebrates database [10]. All but four motifs exhibit zero effect size and thus may be discarded. Of the four non-zero effect-size motifs, MAX::MYC and MYC are nearly identical and one of the pair could be removed.

**Fig S4.**
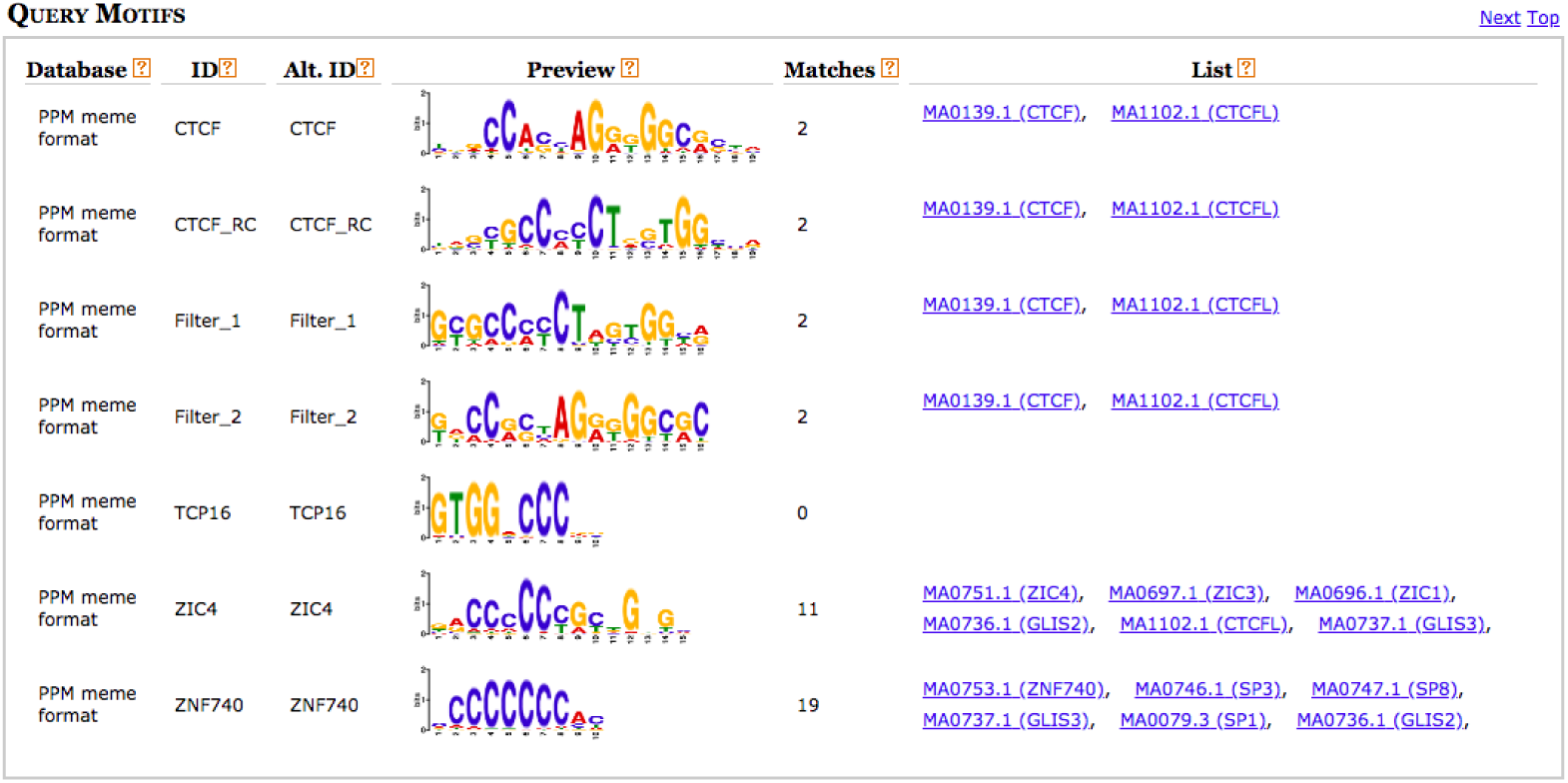
ENCODE Gm12878 CTCF TF ChIP-seq data application Tomtom query. Results from running the Tomtom motif similarity tools [8] querying learned motifs against the 2018 JASPAR Core Non-vertebrates database [10].

## Footnotes

1 The JASPAR naming convention denotes the IRF motif (which we sampled from) as the IRF2 motif. We maintain this distinction throughout the text, i.e. when initializing filters with JASPAR motifs the name IRF2 is used (Supplementary Fig. S3) yet when learning IRF *de novo*, the IRF label is used.

